# D3 dopamine receptors implicate a striatal neuronal subtype in the aversive effects of the antipsychotic drug quetiapine

**DOI:** 10.1101/2025.04.15.649037

**Authors:** Elinor Lewis, Jessie Muir, Ying C. Li, Madeline King, Kaden P. Adams, Julianna Glienke, Sarah Warren Gooding, Kevin J. Bender, Christina K. Kim, Jennifer L. Whistler

## Abstract

Second generation antipsychotics (SGAs) are widely used clinical tools; yet, they often cause negative side effects and take weeks to become effective, leading to poor patient compliance. The effect/side effect profile of individual SGAs is highly variable, and the mechanisms that underlie this variability are not well understood. Here, we identify a role of type 3 dopamine receptor (D3R) neurons in the Nucleus Accumbens (NAc) that mimics the aversive effects of quetiapine. Using single-nucleus RNA sequencing, we show that D3R is expressed in a subpopulation of D1R neurons and defines a distinct NAc cell type. We found that both clozapine and quetiapine cause acute conditioned place aversion in mice, but aversion subsides only after 21 days of treatment with quetiapine, a drug characterized as an arrestin-biased agonist at D3R. We provide evidence at both the cellular and population level that quetiapine inhibits D3R-expressing neurons in the lateral shell (LatSh) of the NAc. We also demonstrate that local injection of quetiapine into LatSh NAc is sufficient to cause place aversion. Selective optogenetic inhibition of D3R-neurons in the LatSh produces real time place aversion in mice, correlating this cell type to the aversive effects of quetiapine. These findings suggest a cell-type-specific mechanism underlying quetiapine-induced aversion and its potential attenuation with chronic treatment, offering insight into the cell types and circuitry that shape the quetiapine side effect profile.

## Introduction

Antipsychotic drugs have become the cornerstone of treatment for schizophrenia^1–3^. First-generation antipsychotics (FGAs), developed in the 1950s, work primarily by strongly blocking dopamine type 2 receptors (D2Rs) and are effective at controlling psychotic symptoms. However, their use is often limited by neurological side effects, particularly extrapyramidal symptoms and tardive dyskinesia^2,3^. In contrast, second-generation antipsychotics (SGAs) were developed in the 1960s with the intention of offering better safety and broader therapeutic benefits^2–5^. SGAs typically show lower affinity to D2Rs and also target a wider range of other G Protein-coupled Receptors (GPCRs)^6^. These differences in pharmacology minimized adverse effects and improve patients’ ability to tolerate long-term treatment, ultimately aiming to reduce the frequency of medication discontinuation^7^.

Today, SGAs are the standard treatment for schizophrenia^8^. Yet SGAs have highly variable effects, and clinicians often use trial and error to identify the most effective treatment for each individual^4,5,8^. Many patients experience intolerable adverse effects, contributing to poor adherence and high discontinuation rates^9^. Despite their pharmacological differences from FGAs, multiple large-scale clinical trials have failed to demonstrate consistent advantages of SGAs in reducing extrapyramidal symptoms or improving overall treatment outcomes^7^. Discontinuation rates for antipsychotic medications remain strikingly high, ranging from 64% to 82%^7,8^. The reasons behind this variability and negative side effects remain unclear, highlighting the need to study SGAs’ mechanism of action in the brain. In addition, SGAs take weeks to reach full efficacy though they likely bind dopamine receptors in the brain within minutes^10,11^, indicating a potential dissociation between the mechanisms underlying the beneficial therapeutic effects and the aversive side effects of these drugs, which often occur immediately.

*In vitro* binding studies indicate that SGAs have equal or higher affinity for the type 3 dopamine receptor (D3R) compared to D2R^12^, yet the functional role of D3R in mediated SGA effects has not been well-characterized. Though sparser than D2Rs, D3Rs are expressed throughout limbic circuits implicated in neuropsychiatric disorders^13–15^. In fact, D3Rs are found in their own distinct population of cortical neurons^16^, as well as in a unique subpopulation of D1R-expressing medium spiny neurons (MSNs) in the Nucleus Accumbens (NAc)^17,18^. Genetic disruption of prefrontal D3Rs evokes reductions in anxiety-like behavior, consistent with therapeutic effects of SGAs^19^, suggesting D3R may be involved in their mechanism of action. Recently, we demonstrated that the ubiquitous SGA quetiapine (Seroquel) functions as arrestin-biased agonist at D3R (antagonist for G protein activity but agonist for arrestin-3 recruitment), leading to functional D3R downregulation and consequent tolerance to the drug’s effects on locomotor activity and calcium channel modulation^20^. These data suggest that D3R may have a key mechanistic role in SGA effects and that variable modulation of arrestin-3 signaling at D3R may influence the therapeutic potential and/or the aversive effects of SGAs. Additionally, attribution of SGA tolerance or side effects to specific cell types could narrow the search for druggable targets and inform the development of better antipsychotic treatments.

In this study, we examined the role of NAc D3R neurons in the aversive effects of SGAs in mice. We have previously demonstrated that the SGAs quetiapine and clozapine differ in their recruitment of arrestin-3 to D3Rs and in their ability to cause functional loss of D3R signaling at the membrane^20^. Building on this, we examined the role of D3Rs in aversion to SGAs. We demonstrate that both clozapine and quetiapine produce conditioned place aversion (CPA) in mice. As clozapine does not cause arrestin-3 recruitment to D3R (antagonist for both G protein activity and arrestin-3 recruitment), this indicates that the D3R-arrestin-3 pathway is not responsible for SGA acute aversive effects. However, we show that mice develop tolerance to the CPA effects of only quetiapine, an arrestin-biased agonist at D3R, and not to clozapine. This tolerance to quetiapine CPA is abolished by blocking arrestin-mediated post-endocytic D3R degradation. These findings led us to investigate the role of D3R-expressing cells in quetiapine-induced aversion and the mechanism through which quetiapine controls their activity. Using *in vivo* calcium imaging and slice electrophysiology, we demonstrate that quetiapine inhibits the activity of D3R-expressing cells in the lateral shell (LatSh) NAc. We further show that optogenetic inhibition of these D3R-neurons is sufficient to produce real time place aversion (RTPA) and that this is specific to LatSh NAc D3R neurons and not medial shell (MedSh) NAc. Our study presents mechanistic insight into the aversive effects of SGAs and suggests potential strategies for isolating these effects while maintaining their therapeutic efficacy.

## Methods

### Animals

All mice were maintained on a 12h reverse light-dark cycle (lights on at 8PM) at 22° C. Mice were group housed with 3-5 same-sex cage mates, given a wheel for enrichment, and had *ad-libitum* access to food and water. All experiments were performed in accordance with guidelines set by the University of California, Davis, University of California, San Francisco, and Princeton University Institutional Animal Care and Use Committees (IACUC).

8-week old C57BL/6J wildtype (WT) mice were purchased from The Jackson Laboratories (#000664). Mice were left undisturbed for at least 1 week after their arrival before the start of experimentation. Transgenic animals (D3-Cre::Flox GASP1^−/0^, D3-Cre::Ai14 (tdTomato)) were genotyped by PCR. D3-Cre::Ai14 (KJ302, MMRRC_034696-UCD) mice were crossed to Flox GASP1^tg/0^ mice (Cyagen) on a C57BL/6 background (see refs ^20,21^). All sample sizes are listed in methods and figure legends. All mice included in this study are males as the D3-Cre transgene is carried on the Y chromosome; it is therefore not possible for us to include females in this genotype.

### Drugs

Quetiapine hemifumarate was purchased from TCI America (111974-72-2, Oregon, USA). Quetiapine was dissolved in 100% dimethyl sulfoxide (DMSO) and then diluted in saline solution to a final concentration of 1.5 mg/ml in 4% DMSO for all systemic injections. For microinjections via fluid cannula, quetiapine was dissolved in 100% dimethyl sulfoxide (DMSO) and then diluted in saline solution to a final concentration of 0.2 or 2 mg/ml in 4% DMSO. Clozapine (Tocris 0444, UK) was dissolved in 100% DMSO, heated at 72° F for 5 minutes, and diluted in saline solution to a final concentration of 0.4 mg/ml in 4% DMSO. Aripiprazole (Sigma SML0935) was dissolved in 100% DMSO heated at 72° F for 5 minutes and diluted in saline solution and

Tween80 to a final concentration of 1.5 mg/ml in 5% DMSO and 10% Tween80. Quetiapine was given to mice at a dose of 15 mg/kg. Clozapine was given at a dose of 4 mg/kg. Aripiprazole was given to mice at a dose of 15 mg/kg. Previous work in our lab shows these doses drive equi-inhibitory effects on locomotion and cocaine-induced locomotion.^20^ We also showed this dose of quetiapine causes tolerance to these locomotor inhibitory effects.^20^ All injections were administered subcutaneously (SC). Vehicle solutions are saline with 4% DMSO or saline to match treatment.

### Statistics

All statistical analyses were performed using R. Test statistics, *n,* and *P* values are provided in figure legends and the supplemental statistics table. Normal distribution and independence within pairs was assumed for paired t-tests, and independence between groups and equal variance were assumed for unpaired t-tests. All outliers were included. Viral expression was confirmed by visualization of dense green fluorescence within the target region as defined by the Allen Mouse Brain Atlas and shown in **Figure S4A.** In D3-Cre::Ai14mice, viral expression specifically in D3 cells was further verified by overlap of green fluorescence with tdTomato-labeled cells. Mice whose post-mortem tissue revealed absent, minimal, or mistargeted expression were excluded. All bars represent mean and error bars represent SEM. Group sizes are based on previous literature^16,18,20^. Investigator was not blinded to experiments. When possible, controls and treatment groups were run side by side with mice from each cage being allocated to either control or treatment groups randomly. Exact statistical parameters and defaults are available at https://github.com/tinakimlab/Lewis-et-al-2025.

### Single Nucleus RNA Sequencing

The medial and lateral NAc of D3-Cre mice (males) were dissected and flash frozen on dry ice. Samples from 2 different mice were pooled for each replicate. Nuclei isolation was performed using the Single Cell 3’ Gene Expression Chromium Nuclei Isolation Kit (10x Genomics, California, USA); cell counts were estimated at 1000 cells/µL using the Countess Cell Counter (Thermofisher, NJ, USA). Barcoded cDNA was generated using the Chromium Single Cell 3’ v3.1 kit, Chromium Next GEM Chip G and Chromium Controller iX (10x Genomics) with a targeted cell recovery of 10,000 cells/µL and libraries were constructed using the Dual Index and Library Construction kits (10x Genomics). Libraries were sequenced using a NovaSeq 6000 (Illumina) at a depth of 300 M reads per sample (see **Supplementary Data)**.

*Analysis of scRNA sequencing data*: Sequencing data were analyzed using the 10X Genomics Cell Ranger 7.10 software and custom-written Python 3 scripts. Briefly, each experimental group was processed separately using CellRanger and subsequently aggregated. We used the 10x provided mouse genome and Cre transcript, including through the 3’ polyA sequence. Resulting data was analyzed using SCANPY (*49*). First, pre-processing was performed to remove the following:

`o` Genes present in fewer than 2 cells.
`o` Cells with fewer than 600 genes
`o` Cells with greater than 6000 genes
`o` Cells with greater than 20 mitochondrial genes

We also removed genes relating to activity and sex (Trf, Plp1, Mog, Mobp, Mfge8, Mbp, Hbb-bs, H2-DMb2, Fos, Jun, Junb, Egr1, Xist, Tsix, Eif2s3y,Uty, Kdm5d) as well as Hbb-bs, H2-DMb2, Fos, Jun, Junb, Egr1, Xist, Tsix, Eif2s3y,Uty, Kdm5d) as well as mitochondrial genes. Data was normalized and log transformed, and highly variable genes were regressed out for clustering. Principal component’s analysis (PCA) was run on the data set; top 26 PCs were used to create the neighborhood graph which was embedded using Uniform Manifold Approximation and Projections (UMAP). The leiden algorithm was used to cluster the neighborhood graph. A summary of the sequencing libraries is provided in Data S2. Top marker genes were used to identify cell types in each cluster by comparing to existing data (data from (*50*)) on Allen Institute). Clusters representing neuronal cell types were isolated and reclustered using the same method described above. Neuronal subtypes were identified using the Allen Institute single cell RNA seq data base. Clusters where Drd1 or Drd3 were found to be significantly overexpressed compared to other clusters were calculated by a Student’s t-test with p-value adjusted for multiple comparisons (*p<0.05, #p<0.1). Matlab scripts are available at https://github.com/tinakimlab/Lewis-et-al-2025.

### Conditioned Place Aversion Assay

Mice were habituated to the CPA boxes (Med Associates Open Field Arena for Mouse ENV-510 in a sound attenuating box, Georgia, USA) for 60 minutes the day before their first recorded session (Day 0). All recordings were captured and analyzed using Med Associates Activity Monitor 7 Software. On every day after habituation, the boxes were outfitted with two distinct contexts: horizontal striped walls, mesh floor, and litsea scent on one side and vertical striped walls, bar floor, and thyme scent on the other. During initial preference day (Day 1), mice had full access to both sides of the box for a 30-minute session. After this, mice were randomly assigned one of the two sides as their drug-paired side. On Day 2-4, mice were given saline and confined to one side of the box for 30 minutes in the morning and given drug and confined to the other side of the box for 30 minutes in the afternoon. Note that on the non-conditioned side, mice were paired with saline instead of vehicle, to avoid giving 4% DMSO to mice two times in one day as required by IACUC. We therefore performed a separate vehicle control, where mice received saline on the non-conditioned side, and 4% DMSO vehicle on the conditioned side. On Day 5 (final preference day), mice were again given access to both sides of the box (no drug present) for 30 minutes, and the time spent on each side was recorded. Data is reported as percent time spent on the drug-paired side on Day 1 versus Day 5. White lights were kept on throughout the recording. All bars represent mean and dots represent individual mice with connecting lines indicating the same mouse at multiple time points. All error bars represent SEM. Significance between baseline and final preference for each group was determined using a Student’s Paired t-test.T-test statistics and group sizes are available in the above table.

### Stereotaxic Surgeries

Mice were anesthetized with 1.5-2% isoflurane and secured with ear bars in the stereotaxic apparatus (RWD, China) on a heating pad. Standard sterile stereotaxic surgery procedures were performed in accordance with approved IACUC protocols. 500nl of virus was injected at 150nl/minute using a Hamilton syringe controlled by an injection pump (World Precision Instruments, Florida, USA). Virus was allowed to diffuse for 5 minutes before withdrawing the needle. Representative histology, including typical viral spread, of MedSh NAc and LatSh NAc targeting is shown in **Figure S4**.

#### Fiber photometry recordings

D3-Cre mice were injected with Syn-Flex-GCaMP6f (Addgene 100833-AAV5). Chronically implantable fibers (Doric Lenses, Quebec, Canada) with a 400 µm core and 0.66 NA threaded through a 1.25 mm diameter stainless steel ferrules were implanted above the viral injection site (LatSh NAc: M/L: 1.9, A/P:2.15, D/V-4.6) to allow for light delivery. Implants were secured to the skull using dental adhesive (Parkell, C&B metabond, New York, USA). Mice were left undisturbed for 5 weeks following surgery to allow sufficient time for viral expression.

#### Optogenetic manipulations

D3-Cre mice were injected with doublefloxed-eNpHR-EYFP (Addgene 20949-AAV9), EF1a-doublefloxed-hChR2(H134R)-EYFP (Addgene 20298-AAV5), or pAAV-Ef1a-DIO-EYFP (Addgene 27056-AAV5). For a subset of eNpHR- or ChR2-control mice, we used mice that had been injected with Syn-Flex-GCaMP6f instead of EF1a-DIO-EYFP. These mice underwent the same stereotaxic procedure in the same brain region and were given blue or orange light in the same manner. WT mice were injected with either CaMKIIa-eNpHR3.0-eYFP (Addgene 26971-AAV1) or CaMKIIa-hChR2(H134R)-EYFP (Addgene 26969-AAV5). A 400µm core, 0.39 NA optical fiber (RWD R-FOC-L400C-39NA) was implanted above the viral injection site (LatSh NAc: M/L: 1.9, A/P:2.15, D/V-4.6, MedSh NAc: M/L: 0.6, A/P:2.15, D/V-4.6). Implants were secured to the skull using dental adhesive (Parkell, C&B metabond). Behavior experiments took place five weeks after stereotaxic surgery.

#### Fluid cannulas

WT mice were implanted with 0.37 NA optical fiber glass fluid cannulas with a 530µm guiding tube and ZF1.25 receptacle (Doric Lenses, Quebec, Canada) (M/L: 1.9, A/P:2.15, D/V-4.6). Implants were secured to the skull using dental adhesive (Parkell, C&B metabond). Behavior experiments took place five weeks after stereotaxic surgery.

### Electrophysiology

Transgenic mice (D3-Cre::Ai14) aged P28-60 were anesthetized under isoflurane. Brains were dissected and placed in 4°C cutting solution consisting of (in mM): 87 NaCl, 25 NaHCO_3_, 25 glucose, 75 sucrose, 2.5 KCl, 1.25 NaH_2_PO_4_, 0.5 CaCl_2_, and 7 MgCl_2_ and bubbled with 5% CO_2_/95% O_2_. 250µm-thick coronal slices that included the ventral striatum were obtained via vibrating blade microtome (VT1200S, Leica, New Jersey, USA). Slices were then incubated for 30 minutes at 33 °C then placed at room temperature until recording. All recordings were performed in ACSF at 32-34°C.

Slices were placed in recording chamber with circulating ACSF perfusion containing (in mM): 125 NaCl, 2.5 KCl, 2 CaCl_2_, 1 MgCl_2_, 25 NaHCO_3_, 1.25 NaH_2_PO_4_, 25 glucose, bubbled with 5% CO_2_/95% O_2_, osmolarity adjusted to ∼300 mOsm. For IPSC recordings, excitatory transmission was blocked with 10µM R-CPP and 10µM DNQX in the ACSF. D3R-expressing neurons were identified tdTomato fluorescence overlaid on a scanning DIC image of the slice. Patch electrodes were pulled from Schott 8250 glass (3-4 MΩ tip resistance). For voltage-clamp IPSC recordings, patch electrodes were filled with an internal solution that contained (in mM): 80 KCl, 40 CsMeSO_4_, 10 HEPES, 4 NaCl, 5 QX-314, 10 Na_2_-phosphocreatine, 4 Mg-ATP, 0.4 Na_2_-GTP, 0.1 mM EGTA; 290 mOsm, pH 7.2-7.25. For drug wash-on, quetiapine (1µM) was bath applied and data was analyzed after 15 minutes of drug application.

Electrophysiological recordings were collected with a Multiclamp 700B amplifier (Molecular Devices, California, USA) and a custom data acquisition program in Igor Pro software (Wavemetrics, Oregon, USA). Voltage-clamp recordings of synaptic currents were acquired at 10 kHz and filtered at 3 kHz. Whole-cell compensation of pipette capacitance and series resistance (R_s_; 50%) were applied upon patch breakthrough. All data were corrected for measured junction potentials of +12 in high Cl^−^ based internal. All recordings were made using a quartz electrode holder to minimize electrode drift within the slice.

Extracellular afferent stimulation was applied through a bipolar silver electrode in theta glass pipette connected to a battery. Pairs of stimuli were elicited through a pulse generator (0.2-0.8 V amplitude, 200 µs duration, 20 Hz frequency) at 15 s intervals to avoid induction of synaptic plasticity. The extracellular stimulating pipette was embedded in the LatSh NAc ∼100 µm lateral from the recorded neuron to avoid direct stimulation. To avoid polysynaptic responses, voltage amplitude/LED power were tuned so that responses were synchronous and monosynaptic and no failures were observed. All bars represent mean and dots represent individual cells with connecting lines indicating the same cell at multiple time points. All error bars represent SEM.

### Fiber Photometry

Fiber photometry recordings were performed using RWD’s Tricolor Multi Channel Fiber Photometry System. Briefly, 470 nm and 410 nm light were alternatively delivered through a 400 μm patchcord (0.57 NA; Doric Lenses) connected to an optical fiber implanted above the NAc. Fluorescence was recorded with a sCMOS camera using RWD software at a frequency of 20 Hz. Animals were head-fixed to prevent patch cord slipping and recorded for 15 minutes both prior to quetiapine injection and 30 minutes after subcutaneous drug injection. Animals were kept on the head fix set-up during the wait period to minimize handling variability. Preprocessing and analysis of photometry data was conducted using custom R scripts https://github.com/tinakimlab/Lewis-et-al-2025). Data was preprocessed using a method adapted from ^22^. Briefly, an adaptive iteratively reweighted penalized least squares (airPLS) algorithm is used to dynamically detect the slope of each channel so it can be subtracted for a flattened trace that maintains high frequency signal changes. A robust linear model was used to predict the standardized 470 nm signal based on the standardized 410 nm signal. First, ΔF/F was calculated by regressing the 410 nm signal onto the 470 nm signal and taking the residuals of this model, which were then divided by the fitted values. The resulting ΔF/F values were then Z score standardized using the median and standard deviation of the baseline period. Finally, a regression model was used to predict the standardized 470 nm signal from the standardized 410 nm signal, and ΔF/F was obtained as the difference between the actual standardized 470 nm values and the predicted values. Peak detection was performed on the standardized z scores by calculating a rolling threshold defined as 2.5 times the median absolute deviation (threshold chosen based on ^23^) of a 21 second rolling window. A custom function was created to ensure peaks are local maxima greater than the absolute threshold of a z score of 2.5. Local maxima that fall below the rolling threshold are considered background signal and excluded. All bars represent mean and dots represent individual mice with connecting lines indicating the same mouse at multiple time points. All error bars represent SEM.

### Microinjections

Mice were anesthetized with 1.5-2% isoflurane and secured with ear bars in the stereotaxic apparatus (RWD, China) on a heating pad. Standard sterile stereotaxic surgery procedures were performed in accordance with approved IACUC protocols. 500 nl of virus was injected at 150 nl/minute using a Hamilton syringe controlled by an injection pump (World Precision Instruments, Florida, USA). Virus was allowed to diffuse for 5 minutes before withdrawing the needle. Mice were then placed in a recovery chamber on a heating pad for 5 minutes. Righting reflex and toe pinch reflex were confirmed, and then mice were placed in the CPP box.

### Histology

Mice were anesthetized under isoflurane and perfused with ice-cold PBS and then with 4% paraformaldehyde. The brain was extracted post-fixed in 4% PFA overnight. The following day, 65-micron slices were cut on a vibratome (Leica). Slices were mounted on slides and imaged on a Keyence BZ-X180 fluorescent microscope.

### Real Time Place Preference Assay

Mice were placed in a box with 2 identical sides, each 1 square foot, with an open door between the two sides. On baseline day, mice were placed in the box for 30 minutes, and their position in the box was recorded. The following day, mice were attached to a patchcord (Doric lenses) and were again placed in the box for 30 minutes. One side of the box was randomly assigned to be the LED side. During this session, each time the mouse entered one side of the box, LED light was delivered through the patchcord connected via a commutator and fiber optic patch cable (400 μm core, 0.39 NA; Thorlabs). For ChR2 mice, blue light (20 Hz, 5 ms pulse width, 5 mW) was delivered. For NpHR mice, constant orange light (595 nm, 5 milliwatts) was delivered when they entered the LED side of the box. Mouse position was tracked and LED was turned on through the camera connected to RWD-OFRS software. Data is displayed as percent time on LED side. Images were analyzed using ImageJ (Fiji). All bars represent mean and dots represent individual mice with connecting lines indicating the same mouse at multiple time points. All error bars represent SEM.

### Quantification of Cell Number

Images of MedSh NAc and LatSh NAc were taken at 40x magnification using the histology methods detailed above. The red channel was used to visualize D3R-tdTomato–expressing neurons, and the blue channel was used to visualize DAPI-stained nuclei. Cell bodies were initially segmented using Cellpose (cellpose.org), and additional masks were manually added as needed. Mask sizes and positions were measured using ImageJ (Fiji). Raw mask data are provided in the supplemental materials. Bars represent mean and dots represent number of cells in the area from an individual mouse. All error bars represent SEM.

### Code and Data Availability

All R and Matlab scripts used for analysis can be found at https://github.com/tinakimlab/Lewis-et-al-2025. RNA sequencing raw data is available at Ihttps://www.ncbi.nlm.nih.gov/geo/ (GEO Accession GSE329758), and processed RNA sequencing data is available in Supplemental data files. All raw data is available on FigShare (https://doi.org/10.6084/m9.figshare.30906632).

## Results

### Single nucleus RNA sequencing reveals D3-receptor expressing cell-subtype

To begin to assess the role of D3Rs in SGA effects/side effects, we determined the expression patterns of D3R across cell-types in the NAc. NAc from transgenic mice expressing Cre in D3 cells (D3-Cre) were micro-dissected and dissociated to generate a barcoded single nucleus RNA library for next-generation sequencing. Dimensionality reduction and unsupervised clustering of the sequencing data identified the expected main cell-types present in the NAc. (**Figure S1**). Neuronal cell-types were isolated and reclustered for further analysis (**Figure 1A**).

**Figure 1:**
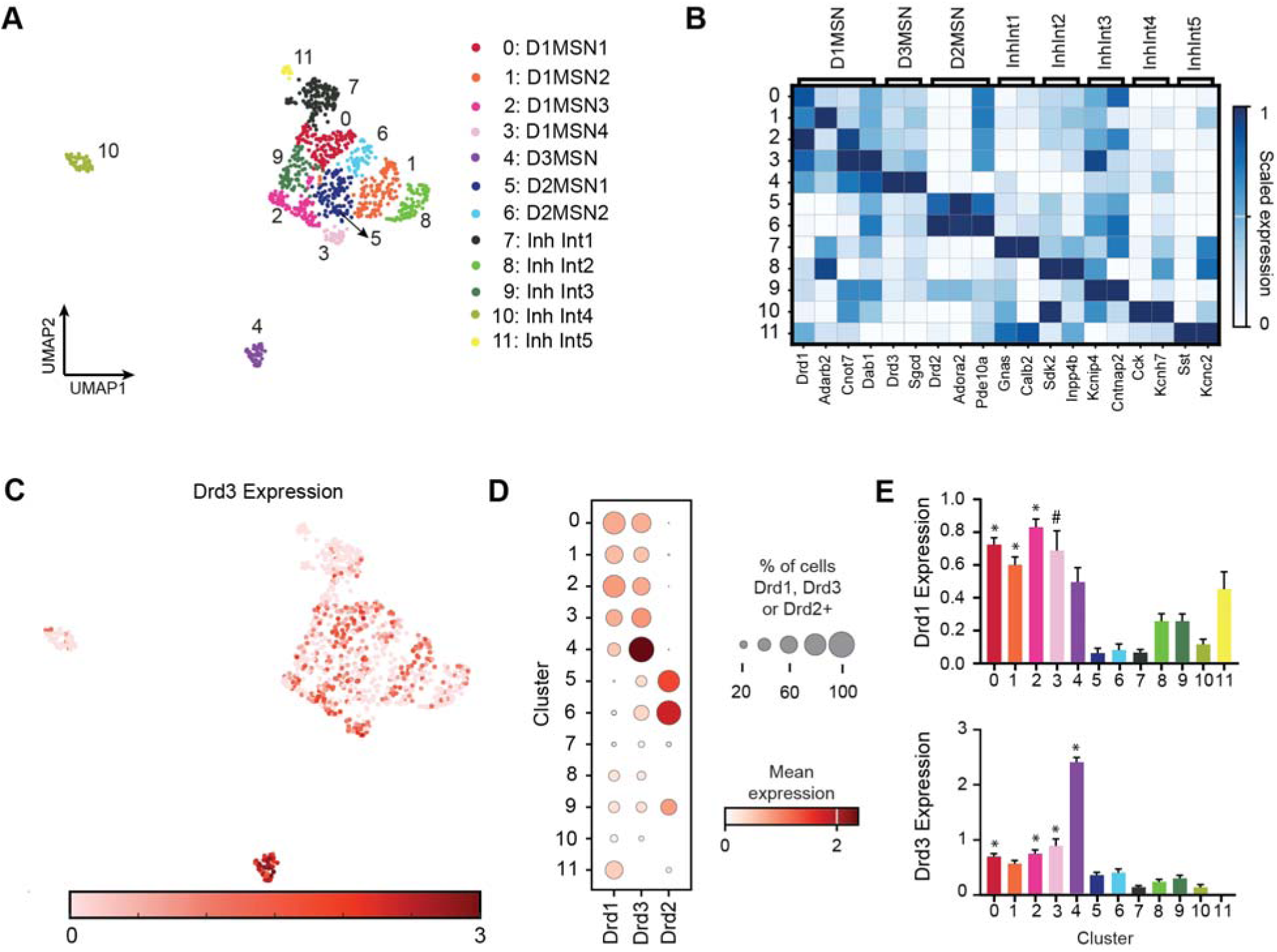
snRNA-seq of NAc neurons reveals D3R expressing cell subtype. A) UMAP visualization of individual nuclei corresponding to different neuronal cell-types. B) Heatmap showing the expression of representative cell-type marker genes reported from prior mouse NAc RNA sequencing datasets. C) UMAP visualization of log1p normalized D3R expression in all nuclei across clusters. D) Dot plot representing the fraction of nuclei within each cluster that are Drd1, Drd2 or Drd3+ (unit variance scaled, log1p normalized count > 0). E) Mean log1p normalized Drd1 and Drd3 in each cluster. * Denotes clusters where Drd1 or Drd3 were found to be significantly overexpressed compared to other clusters, calculated by a Student’s t-test with p-value adjusted for multiple comparisons (*p<0.05, #p<0.1). Error bars represent SEM. See Supplemental Data S1 for additional statistics.

We identified 12 unique subclusters of neurons distinguished based on their expression of marker genes consistent with previous literature including 4 subtypes of D1-MSN, 2 subtypes of D2-MSN, and several types of inhibitory interneurons (**Figure 1B**)^24–26^. Drd3 (D3R gene) expression was enriched in several subclusters (**Figure 1C-E**). Consistent with previous literature, Drd3 is enriched in a subcluster of D1-MSNs (Cluster 2) (**Figure 1D, E**), and chi-squared analysis reveals that the number of cells co-expressing Drd3 and Drd1 significantly outnumbered those co-expressing Drd3 and Drd2. Interestingly, we identified one cluster of cells (labeled D3MSN) that showed greater enrichment of Drd3 compared to Drd1 (**Figure 1D, E**). These data indicate that D3R expression is not restricted to predominantly D1-MSNs but is enriched in a unique cell subtype.

### D3R-GASP1 Trafficking Modulates Tolerance to Quetiapine-Induced Aversion

SGAs can induce side effects such as dysphoria^9,17,27^, thought to involve the NAc^3^. Given that SGAs bind to D3Rs and some SGA variability can be attributed to distinct signaling from D3R^20^, we hypothesized that D3Rs in the NAc may contribute to this aversive response. To model acute aversive side effects of SGAs in mice, we used a conditioned place avoidance (CPA) paradigm (**Figure 2A**). Mice were placed in boxes with two distinct sides, each characterized by unique wallpaper, flooring, and scent. On the day of initial preference testing, mice could freely explore both sides. For the next three days, they were confined to one side of the box for 30 minutes in the morning following a saline injection and then confined to the opposite side in the afternoon after injections of either vehicle (4% DMSO), clozapine (4 mg/kg), or quetiapine (15 mg/kg). No injections were given on the final preference testing day on which mice were given free access to both sides of the chamber. Time spent on each side was assessed relative to their recorded baseline preference prior to the drug conditioning paradigm. Mice showed no preference or aversion for the vehicle-treated side compared to saline (**Figure 2B**) but showed place aversion to the context associated with either clozapine or quetiapine (**Figure 2C, D**).

**Figure 2:**
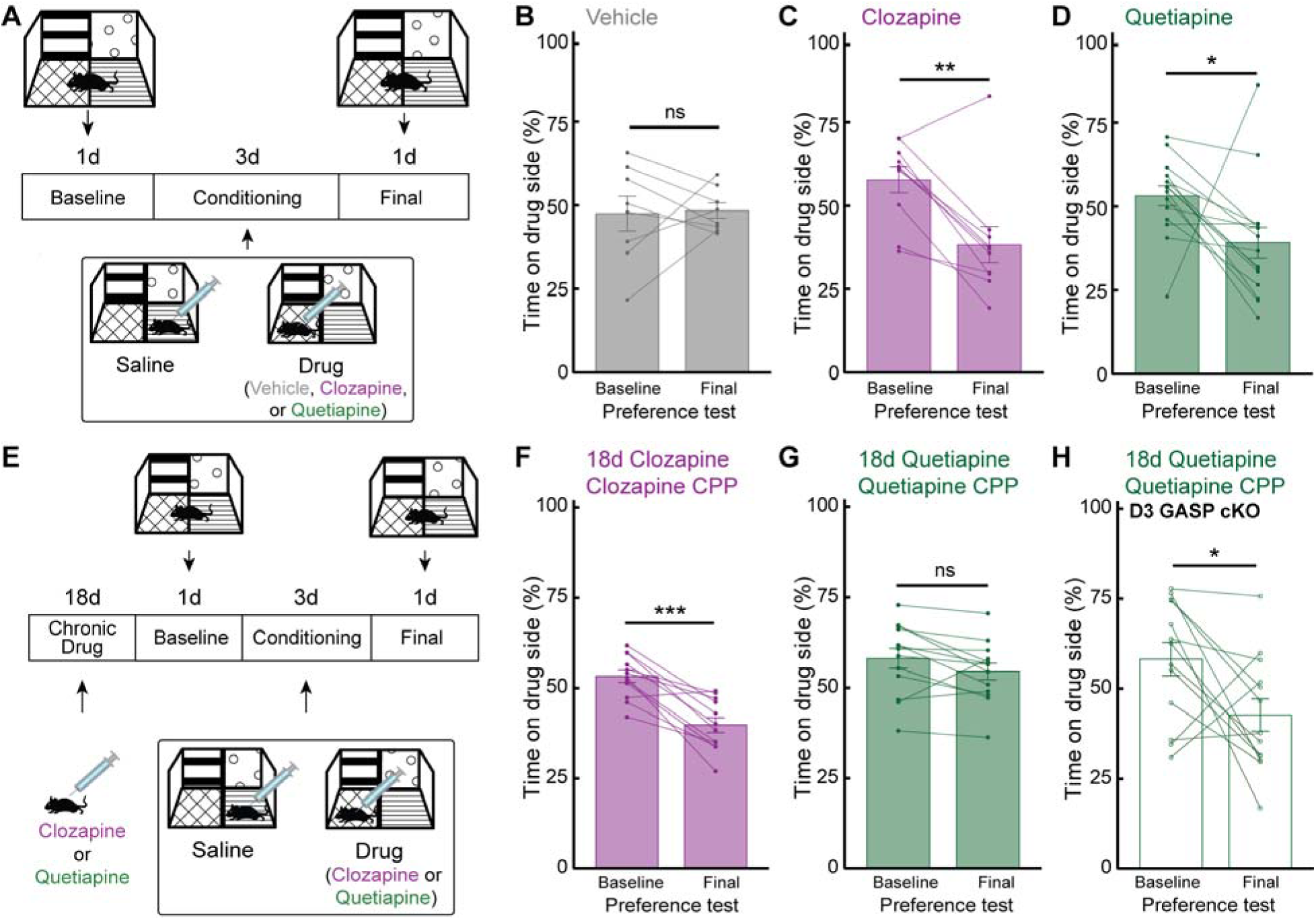
Mice become tolerant to the aversive effects of quetiapine but not clozapine. A) Timeline of conditioned place experiment. Baseline preference test and final preference test take place when mice are off drug. Conditioning period includes AM saline treatment in one context and PM drug treatment in the other context (4% DMSO vehicle, 4 mg/kg clozapine, or 15 mg/kg quetiapine s.c.). B) Time spent on vehicle side before and after conditioning (N=8 mice, Student’s Paired t-test, ns p=0.868, t=-0.172). C) Time spent on clozapine side before and after conditioning (N= 10 mice, Student’s Paired t-test, *p=0.002, t=4.251). D) Time spent on quetiapine side before and after conditioning (N=15 mice, Student’s Paired t-test, *p=0.044, t=2.215). E) Timeline of chronic treatment experiment. Mice were treated with clozapine (4 mg/kg) or quetiapine (15 mg/kg) daily s.c. and then place conditioned as in A). F) Time spent on clozapine side before and after conditioning following chronic clozapine (N=12 mice, Student’s Paired t-test, ***p=7.72e-05, t=6.102). G) Time spent on quetiapine side before and after conditioning following chronic quetiapine (N=13, Student’s Paired t-test, ns p=0.063, t=2.052). H) Time spent on quetiapine side after conditioning following chronic quetiapine in D3 GASP cKO mice (N=13 mice, Student’s Paired t-test, *p=0.032, t=2.430). All error bars represent SEM.

Both clozapine and quetiapine are D3R G protein antagonists but only quetiapine promotes arrestin-3-mediated loss of D3R function^20^. To interrogate the role of D3Rs in SGA-induced CPA, we assessed whether chronic treatment with quetiapine or clozapine altered place aversion (**Figure 2E**). Mice were treated for 18 days with drug before undergoing the CPA paradigm (21 days total drug). Chronic treatment with clozapine had no effect on clozapine CPA as clozapine pre-treated mice retained an aversion to the drug paired side (**Figure 2F**). However, chronic quetiapine treatment abolished quetiapine-induced CPA (**Figure 2G**). Chronic vehicle treatment had no effect on clozapine or quetiapine CPA, as expected (**Figure S2A-C**).

We have previously shown that functional downregulation of D3Rs in response to chronic quetiapine requires arrestin-mediated endocytosis and post-endocytic D3R degradation through its interaction with the GPCR-associated sorting protein-1 (GASP1)^20,28^. We hypothesized that this mechanism also mediated tolerance to quetiapine. To examine this, we repeated the CPA experiments in transgenic mice with a selective deletion of GASP1 only in D3R-expressing cells (D3 GASP1 cKO). We found that chronic quetiapine treatment does not produce tolerance to quetiapine-induced CPA in D3 GASP1 cKO mice (**Figure 2H**). These data suggest that D3R signaling plays a role in quetiapine aversion and that tolerance to quetiapine’s aversive effects may involve GASP1-mediated functional downregulation of D3Rs. To examine whether acute clozapine aversion is also mediated through D3R, we performed a cross-tolerance experiment and examine the effects of chronic quetiapine treatment on clozapine CPA. We found that quetiapine pre-treatment did not abolish clozapine CPA (**Figure S2D, E**), indicating loss of D3Rs is not sufficient to eliminate the aversive effects of clozapine. To examine whether another arrestin-biased agonist at D3R caused the same effect as quetiapine, we performed an acute CPA and subsequent tolerance to CPA experiment with aripiprazole (Abilify) and saw aversion to aripiprazole and the abolishment of this aversion after chronic treatment **(Figure S5),** consistent with a role for D3R in this effect, though aripiprazole’s additional partial agonism at D2R may also contribute to driving aversion^29^.

### Quetiapine drives inhibition of D3R neurons in the LatSh NAc

We next interrogated the mechanism by which quetiapine altered the activity of D3R-expressing neurons in the NAc. Quetiapine has been shown to modulate T-type calcium channels at the axon initial segment of layer V pyramidal neurons in the prefrontal cortex (PFC) via D3R and arrestin-3-dependent pathways^16,20,30^. However, quetiapine’s impact on neuronal activity in midbrain regions critical for CPA has not been investigated^31–33^. A subtype of D1 MSNs in LatSh NAc has recently been implicated in aversion to foot shock^18^, providing a candidate subpopulation of D3R cells mediating SGA aversion. We first examined the effects of quetiapine on D3R neuronal activity *in vivo*, using a genetically encoded fluorescent calcium sensor (flex-GCaMP6f) to monitor activity with fiber photometry (**Figure 3A**). We found that, 30 minutes following a systemic quetiapine injection, D3R neurons exhibit reduced frequency of calcium transients compared to the baseline period, while a vehicle injection elicited no change (**Figure 3B-D**). We next examined the mechanism by which these decreases in activity occur using whole cell patch clamp electrophysiology (**Figure 3E**). To visualize D3R-expressing neurons, we crossed Ai14 reporter mice which express a Cre-dependent tdTomato reporter to D3R-Cre mice. We found that quetiapine increases the amplitude of evoked inhibitory post synaptic currents (IPSCs) with no subsequent change in paired pulse ratio or coefficient of variation, consistent with a postsynaptic effect (**Figure 3F-H**). Together, these *ex vivo* and *in vivo* data suggest that quetiapine inhibits D3R-expressing neurons in the LatSh NAc.

**Figure 3:**
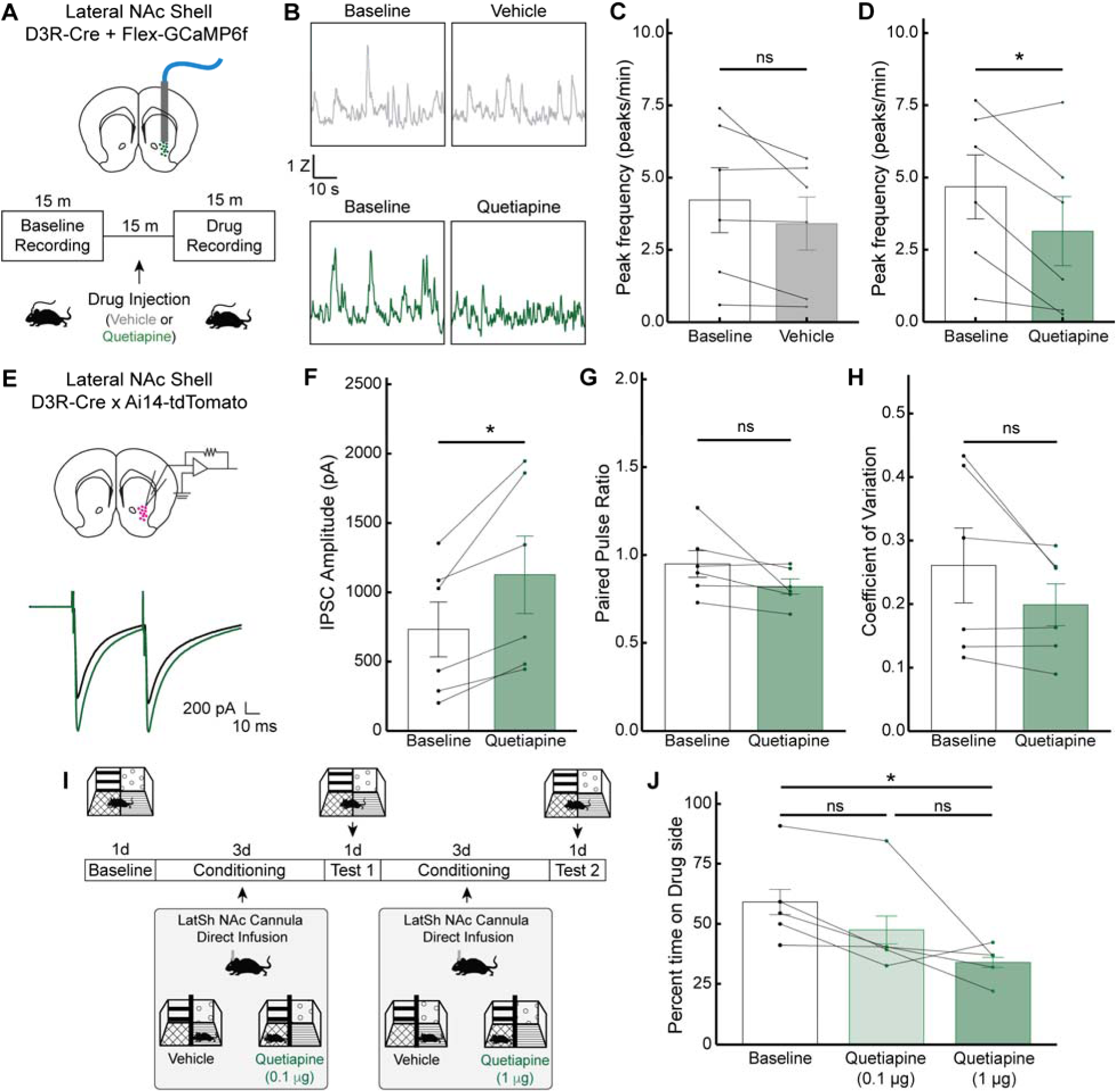
Quetiapine causes inhibition of D3R neurons in LatSh NAc. A) Schematic and timeline of fiber photometry experiment. A 15 minute baseline recording was taken of GCaMP6f activity in LatSh NAc D3R neurons. Mice were injected s.c. with drug or 4% DMSO vehicle and then recorded again 15 minutes later for a 15 minute drug recording. B) Sample traces of GCaMP6f recordings from LatSh NAc D3R neurons. C) Mean GCaMP6f peak frequency before and after vehicle injection (N=6 mice, Student’s Paired T-test, t=0.422, ns p=0.122). D) GCaMP6f peak frequency before and after quetiapine injection (N=6 mice, Student’s Paired T-test, t=3.0571, *p=0.038). E) Example current traces evoked by paired-pulse synaptic stimulation before (black) and after (green) application of quetiapine (1 µM) from D3R tdTomato+ neurons in LatSh NAc. Values shown in F-H are average from 20 traces over 5 minutes. F) Amplitude of evoked IPSCs before and after quetiapine (N=6 cells, Student’s Paired T-test, t=3.687, *p=0.014). G) Paired pulse ratio before and after quetiapine (N=6 cells, Student’s Paired T-test, ns, p=0.140, t=1.754). H) Coefficient of variation before and after quetiapine (N=6 cells, Student’s Paired T-test, ns, p=0.129, t=1.818). I) Timeline of conditioned place experiment with cannula direct infusion of quetiapine. Baseline preference test and final preference test take place when mice are off drug. Conditioning period includes AM saline treatment in one context and PM drug treatment in the other context. See methods section for more detail. J) Time spent on drug paired side after conditioning (one-way repeated-measures ANOVA with Tukey post-hoc tests; N=5 mice; main effect: p=0.037, F_2,8_=5.11; baseline vs 1 µg: *p=0.031, t=3.195; baseline vs 0.1 µg: ns, p=0.349, t=1.481; 0.1 µg vs 1 µg: ns, p=0.258, t=1.713). All error bars represent SEM.

To assess whether quetiapine could be mediating its aversive behavioral effects through mechanisms of action in the LatSh NAc, we then repeated the CPA paradigm in Figure 2 with direct injections of quetiapine into LatSh NAc using a microinjection fluid cannula (**Figure 3I**). We found a dose-dependent effect of drug conditioning, with mice exhibiting place aversion to the side of the box that had been paired with a 1µg direct injection of quetiapine into the LatSh NAc (**Figure 3J**).

### Optogenetic inhibition of D3R neurons in the LatSh but not MedSh NAc drives aversion

We sought to determine whether inhibition of LatSh D3R-expressing neurons is sufficient to mimic the aversive effects of locally administered quetiapine in the LatSh. To accomplish this, we injected a Cre-dependent halorhodopsin (eNpHR, an orange light-activated chloride pump) in the LatSh of D3R-Cre mice to selectively express the inhibitory opsin in D3R neurons (eNpHR+; **Figure 4A**). This approach allowed us to selectively inhibit LatSh D3R cells during a Real-Time Place Aversion (RTPA) paradigm. Control mice did not express eNpHR but were still given orange light stimulation in LatSh NAc (eNpHR-). On the baseline day, mice were placed in a box with two identical sides and allowed to move freely between sides. The following day, mice returned to the arena but entry into one side was paired with LED illumination to activate eNpHR, thus inhibiting activity of LatSh NAc D3R neurons whenever they occupied that side of the box (**Figure 4B**). We found that mice expressing eNpHR in D3R neurons avoided the light-paired side, whereas the control mice showed no change in preference to the orange light-paired side (**Figure 4C**). The fold change in preference for the light-paired side was also significantly lower in eNpHR+ compared to eNpHR- mice (**Figure 4D**).

**Figure 4:**
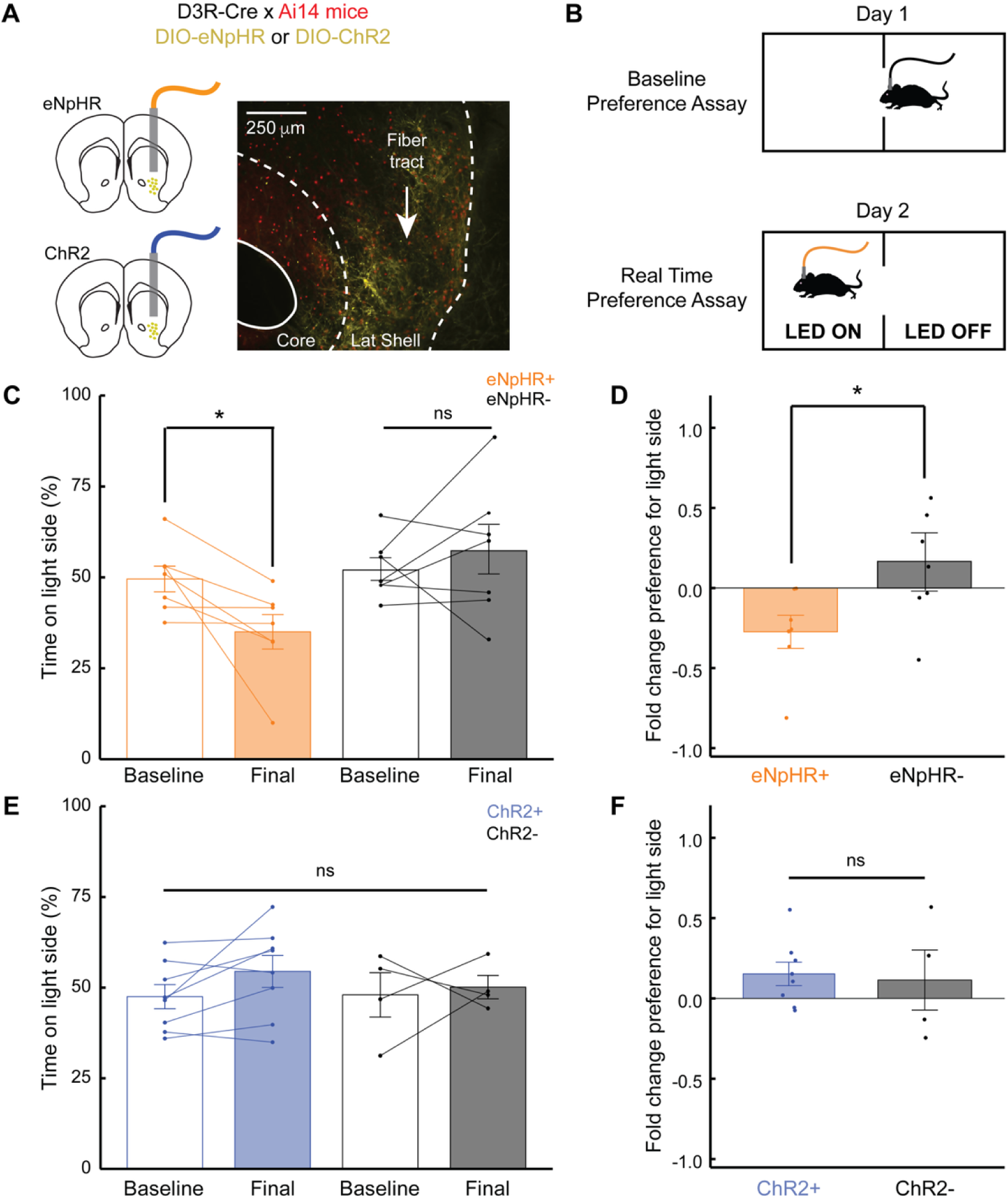
Optogenetic inhibition of D3R neurons in LatSh NAc produces real time place aversion in mice. A) Schematic and histology. D3R-Cre x Ai14-tdTomato mice were injected with either AAV-EF1a-DIO-eNpHR-eYFP (eNpHR+) or AAV-EF1a-DIO-ChR2-eYFP (ChR2+) in LatSh NAc. Control mice did not express an opsin in LatSh NAc but were otherwise treated the same (eNphR- or ChR2-). B) Schematic of Real Time Place Assay. LED is orange for eNpHR mice and blue for ChR2 mice. C) Time spent on light side during baseline versus final test for eNpHR+ and eNpHR- mice (Two-way ANOVA with Tukey post-hoc tests; N=7,7 mice; interaction: *p=0.048, F_1,12_ =4.855; eNpHR+ baseline vs final: *p=0.040, t=2.299; eNpHR- baseline vs final: ns, p=0.430, t=-0.818). D) Fold change from baseline preference to real time preference for orange light side in eNpHR+ compared to eNpHR- mice (Welch unpaired T-test, * p=0.003, t=-3.327). E) Time spent on light side during baseline versus final test for ChR2+ and ChR2- mice (Two-way ANOVA; N=8,4 mice; interaction: ns, p=0.516, F_1,10_=0.453). F) Fold change from baseline preference to real time preference for blue light side in ChR2+ mice compared to ChR2- mice (Welch unpaired T-test, ns, p=0.782, t=0.285). All error bars represent SEM.

We repeated this paradigm using a Cre-dependent channelrhodopsin (ChR2, a blue light-activated cation channel) to determine the behavioral effect of activating this subpopulation. Neither ChR2+ nor ChR2- mice exhibited any preference for the blue light-paired side (**Figure 4E**), and there was no difference in the fold change preference for the light-paired side across days between ChR2+ and ChR2- mice (**Figure 4F**). In contrast, wild-type (WT) mice controls expressing a nonspecific eNpHR in LatSh NAc neurons showed no preference or aversion to the LED side (**Figure S3A**), while WT mice expressing nonspecific ChR2 in LatSh NAc neurons showed a clear preference for the LED illuminated side, (**Figure S3B**). These data suggest that the real-time aversion phenotype is driven by inhibition of the D3R population specifically within the LatSh.

Given that D3R neurons are also strongly present in the MedSh NAc (**Figure S4A, B**), we also examined the effects of quetiapine and D3R inhibition in this subregion. We performed GCaMP6f fiber photometry in D3R neurons in MedSh NAc during systemic quetiapine injection and did not observe any robust changes in calcium transient frequency after vehicle or quetiapine injection (**Figure 5A-D**). We then expressed either eNpHR or ChR2 in D3R neurons in MedSh NAc and performed the RTPA during inhibition or excitation of these neurons, respectively (**Figure 5E, F**). Control mice did not express eNpHR or ChR2 but were still given blue light in MedSh NAc. Neither eNpHR-mediated inhibition (**Figure 5G, H**) nor ChR2-mediated activation (**Figure 5I, J**) of the D3R-expressing MedSh NAc neurons produced an effect on RTPA. Histology quantification in D3R-Cre x Ai14-tdTomato mice confirmed that there are significantly more D3R+ neurons in the MedSh compared to the LatSh, ruling out the possibility that the null behavioral results with MedSh perturbation were due to a lack of D3R+ neurons in the MedSh NAc (**Figure S4C, D**).

**Figure 5:**
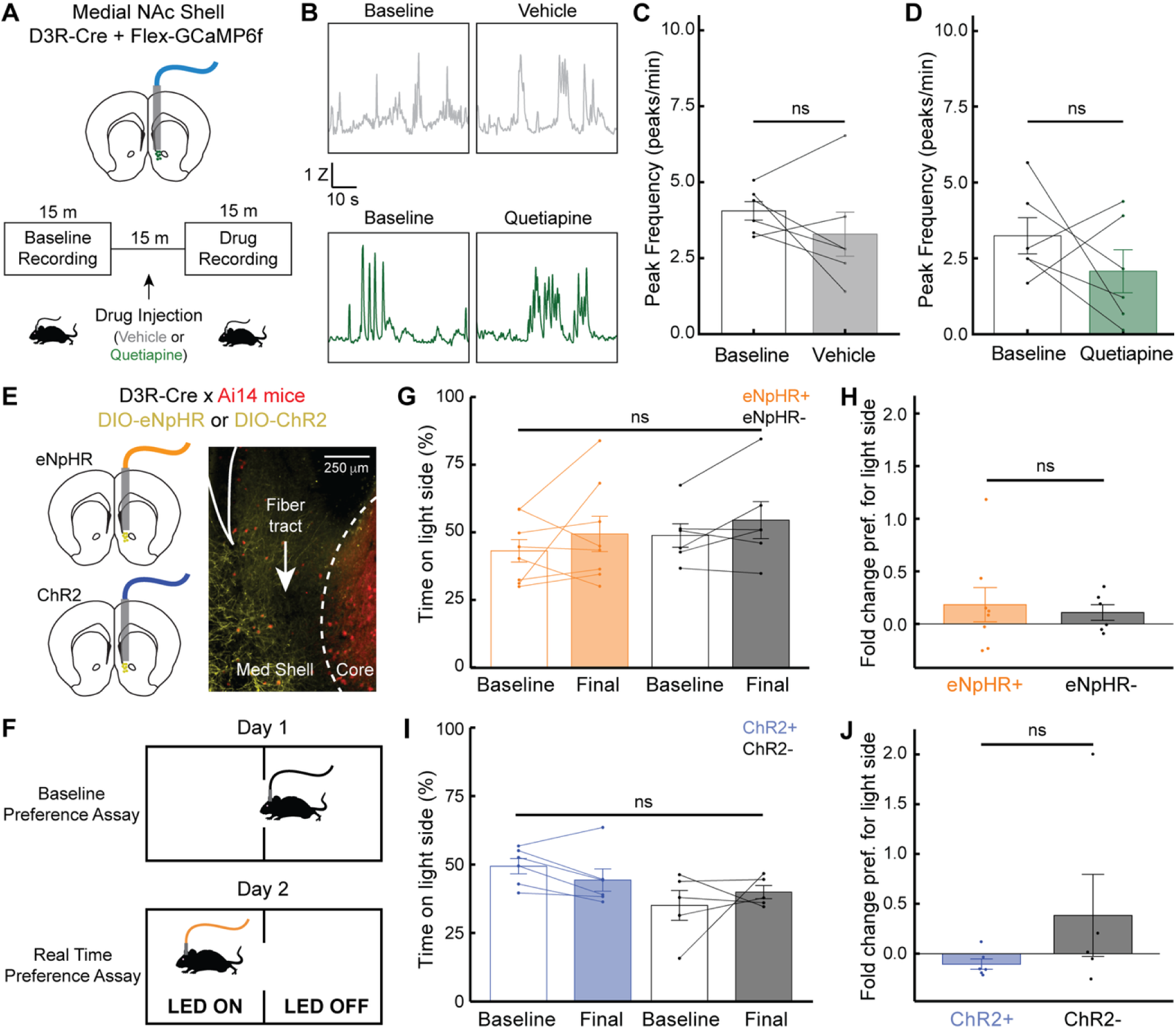
Neither Inhibition nor activation of D3R neurons in MedSh NAc produces real time place aversion. A) Schematic and timeline of fiber photometry experiment. A 15 minute baseline recording was taken of GCaMP6f activity in MedSh NAc D3R neurons. Mice were injected s.c. with drug or 4% DMSO vehicle and then recorded again 15 minutes later during a 15 minute drug recording. B) Sample traces of GCaMP6f recordings from MedSh NAc D3R neurons. C) Mean GCaMP6f peak frequency before and after vehicle (N=6 mice, Student’s Paired T-test, t=1.133, ns p=0.309). D) Mean GCaMP6f peak frequency before and after quetiapine (N=6 mice, Student’s Paired T-test, t=1.068, * p=0.334). E) Schematic and histology. D3R-Cre x Ai14-tdTomato mice were injected with either AAV-EF1a-DIO-eNpHR-eYFP (eNpHR+) or AAV-EF1a-DIO-ChR2-eYFP (ChR2+) in MedSh NAc. Control mice did not express an opsin in MedSh NAc but were otherwise treated the same (eNphR- or ChR2-). F) Schematic of Real Time Place Assay. LED is orange for eNpHR mice and blue for ChR2 mice. G) Time spent on light side during baseline versus final test for eNpHR+ and eNpHR- mice (Two-way ANOVA; N=8,6 mice; interaction: ns, p=0.943, F_1,12_=0.0053). H) Fold change from baseline preference to real time preference for orange light side in eNpHR+ compared to eNpHR- mice (Welch unpaired T-test, ns, p = 0.110, t=0.610). I) Time spent on light side during baseline versus final test for ChR2+ and ChR2- mice (Two-way ANOVA; N = 6,5 mice; interaction: ns, p=0.196, F_1,9_=1.96). J) Fold change from baseline preference to real time preference for blue light side in ChR2+ compared to ChR2- mice (Welch unpaired T-test, ns, p=0.548, t=-1.77). All error bars represent SEM.

Together these results indicate that inhibition of D3R expressing neurons specifically in the LatSh NAc drives the aversive behavioral effects seen with both systemic and local quetiapine injections in mice.

## Discussion

SGAs are widely prescribed to manage severe mental illness (SMI)^1,5,6^. Despite their critical role in treatment, the mechanisms underlying their therapeutic effects remain poorly understood. Many SGAs were developed before the discovery of D3R^34^, warranting additional study in the context of this receptor. While SGAs are critical for treatment of SMI, they often cause adverse side effects that compromise patient compliance^7^. In this study, we explored the role of D3R in SGA-induced aversion using quetiapine and clozapine, two commonly prescribed SGAs that are both full antagonists at D2R^29,35^ but have differing abilities to recruit arrestin-3 to D3R^20^ and drive functional downregulation of D3Rs^20^. We note that future studies are needed to examine how other SGAs modulate LatSh NAc activity and aversion. Notably, aripiprazole — which, like quetiapine, acts as an arrestin-biased agonist at D3R^20^ — also produces acute CPA and tolerance to CPA with chronic pre-treatment **(Figure S5),** consistent with a role for D3R in this effect. However, as aripiprazole is also a partial agonist at D2R^29^, its D2R activity cannot be ruled out as a contributing factor, and further mechanistic dissection of aripiprazole tolerance remains an important direction for future work. Additional SGAs such as cariprazine, which has high affinity for both D2R and D3R but acts as a partial G protein agonist rather than an antagonist, also warrant future investigation. As we did not examine the effects of FGAs on LatSh NAc D3R activity, additional experiments are needed to determine whether the modulation of D3Rs shown here is unique to SGAs.

In addition, it is well known that rodent models of psychosis fall short in capturing the complex and heterogeneous underlying biology of schizophrenia. Furthermore, our behavioral measurements of aversion in mice, while robust and reproducible, may not represent the full spectrum of aversive side effects that can be experienced in patients. Nonetheless, studying how SGA-induced circuit level changes drive behavioral aversion in rodents is a critical step in understanding the mechanism of action of these drugs in the context of the mammalian brain. The ability to expand beyond profiling the pharmacology of SGAs in culture, to study their effects on cell-type-specific neuronal activity and aversive behaviors, is a critical step towards eventually improving their use in patients.

Both clozapine and quetiapine produce acute aversive effects in mice; however, only quetiapine promotes tolerance to this aversion over time. This tolerance to quetiapine is dependent on the post-endocytic sorting protein GASP1 in D3R neurons (**Figure 2H**), implicating post-endocytic trafficking of D3Rs (and not effects at D2R) in tolerance. Supporting our findings, prior studies have suggested that neither quetiapine nor clozapine promotes arrestin recruitment at D2Rs^29,35^. However, there are currently no tools sufficient for selectively quantifying D3R number independently of D2R number^15,36^, and thus potential chronic effects of quetiapine, specifically at D2R, cannot be ruled out. Thus future studies are needed to determine whether additional receptor targets may also play a role in tolerance to quetiapine’s aversive effects.

Mechanistically, we found that D3R neurons in the LatSh NAc show reduced activity in response to quetiapine injection *in vivo*, and these neurons also show increased evoked IPSC amplitude *ex vivo* (**Figure 3A-H**), consistent with a postsynaptic inhibitory mechanism^37,38^, though the precise downstream signaling remains an important open question. To tie this inhibition to the aversive effects of quetiapine, we found that local injection of quetiapine into LatSh NAc produced CPA (**Figure 3J**) and that specific optogenetic inhibition of D3R neurons in the LatSh NAc produced RTPA (**Figure 4**). These findings suggest that D3R-expressing neurons in the LatSh NAc (but not MedSh) play a role in mediating aversive effects of quetiapine and that clozapine produces aversion through additional mechanisms. Future studies can examine whether D3R neurons in brain regions outside of the NAc are similarly inhibited by quetiapine, and whether this inhibition can drive aversion.

Medium spiny neurons (MSNs) in the NAc are typically classified into two major genetic subpopulations based on dopamine receptor expression: D1R-expressing and D2R-expressing neurons^39,40^. However, numerous genetically distinct subpopulations are contained within each of these groups, and our sequencing data indicate that D3Rs also define a unique population of neurons (**Figure 1A**). One limitation of our data is that the tissue used for RNA sequencing was obtained from the entire LatSh NAc region using anatomically-guided microdissection; thus future studies using spatial transcriptomics are needed to delineate the precise subregional location of these distinct cell-type clusters throughout the LatSh NAc area.

Given that several, but not all, subtypes of D1R-neurons within the LatSh NAc have been identified as a source of RTPA^18,41^ and that D3R is strongly co-expressed with D1R in the NAc (**Figure 1D**), we hypothesized that D3R-expressing neurons might represent a key subpopulation mediating aversion. *In vivo* neural recordings revealed inhibition following quetiapine injection of D3R-expressing cells, while slice electrophysiology recordings suggest this is driven by an increased IPSC amplitude in these neurons (**Figure 2**). Similarly, optogenetic inhibition of D3R-expressing neurons specifically in the LatSh NAc drove an aversive response. One limitation of this study is that we cannot rule out that quetiapine is altering activity of D3R-neurons by binding to receptors other than the D3R. That said, it is unlikely this is through binding to D2R, which is structurally similar to D3R, as D2R and D3R are expressed in distinct populations of MSNs (**Figure 1D, E**).

The medial and lateral shell regions of the NAc are often described as having opposing roles, with the MedSh region more commonly associated with reward and the LatSh with aversion^42–46^. Increasing evidence demonstrates these labels are oversimplistic, and that both reward and aversion are shaped by more nuanced, projection-specific, and genetically distinct neuronal subpopulations within each region^47^. In this study, while inhibition of D3R LatSh neurons drove a real-time place avoidance, nonspecific inhibition of all LatSh NAc neurons did not (**Figure S3A**). Furthermore, inhibition of D3R neurons in the MedSh NAc did not drive a real-time place avoidance either (**Figure 5G**). Given the heterogeneity in effects of VTA dopamine modulation in different subdivisions of the NAc (MedSh, LatSh, and core) during reward and aversion^47–53^, it is likely the D3R LatSh NAc’s contribution to aversion is driven by neuronal subpopulations with distinct downstream connectivity, rather than aversion being a uniform function of D3R neurons across the entire NAc. Our studies confirm that aversion can be mediated by a specific D3R-expressing population in the LatSh.

Our findings reveal a model of aversion to quetiapine spanning cellular and system levels and demonstrate selective tolerance to this aversion based on receptor dynamics. The D3R cells in the NAc LatSh are a unique subpopulation and may express other potentially druggable targets not expressed in other NAc neurons. Our findings therefore could provide a framework to design future therapeutics that maximize the therapeutic efficacy of SGAs while reducing the side effects that limit compliance.

## Author Contributions

EL, CKK, JLW, and KJB conceived the project. EL, JM, YL, SWG, KJB, CKK, and JLW contributed to data analysis. EL, JM, YL, KPA, MK, and JG contributed to data collection. EL was responsible for the original draft. KJB, CKK, and JLW supervised the project and acquired funding.

---

**Figure S1:**
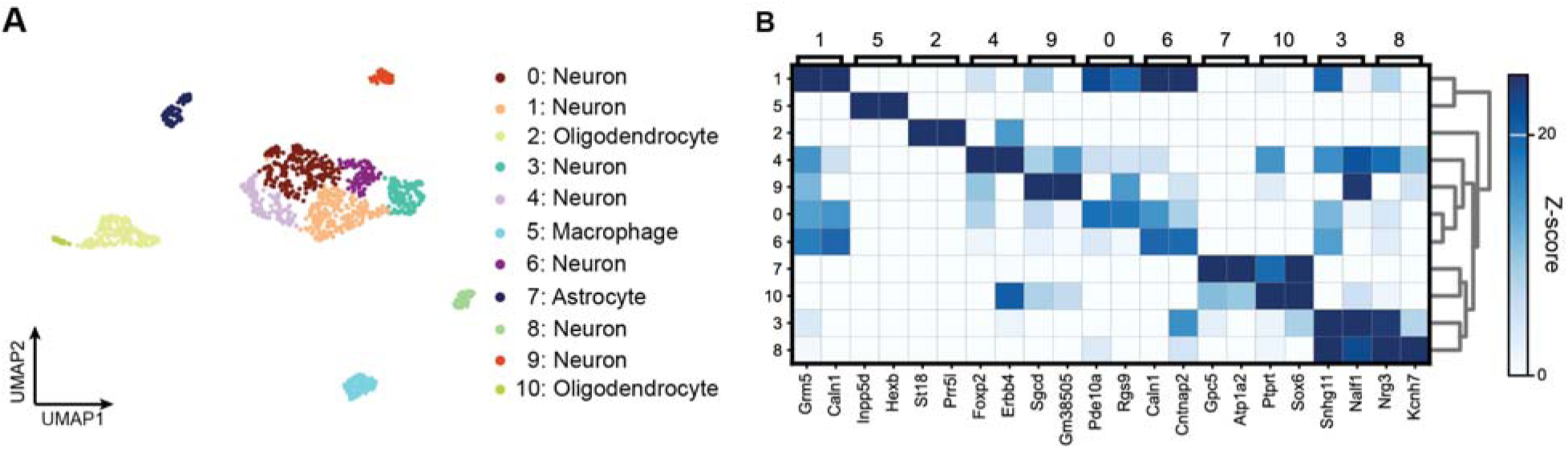
Additional snRNA-seq analysis. A) UMAP visualization of individual nuclei corresponding to different cell-types. B) Heatmap showing the expression of representative cell-type marker genes. See Supplemental Data S1 for additional statistics.

**Figure S2:**
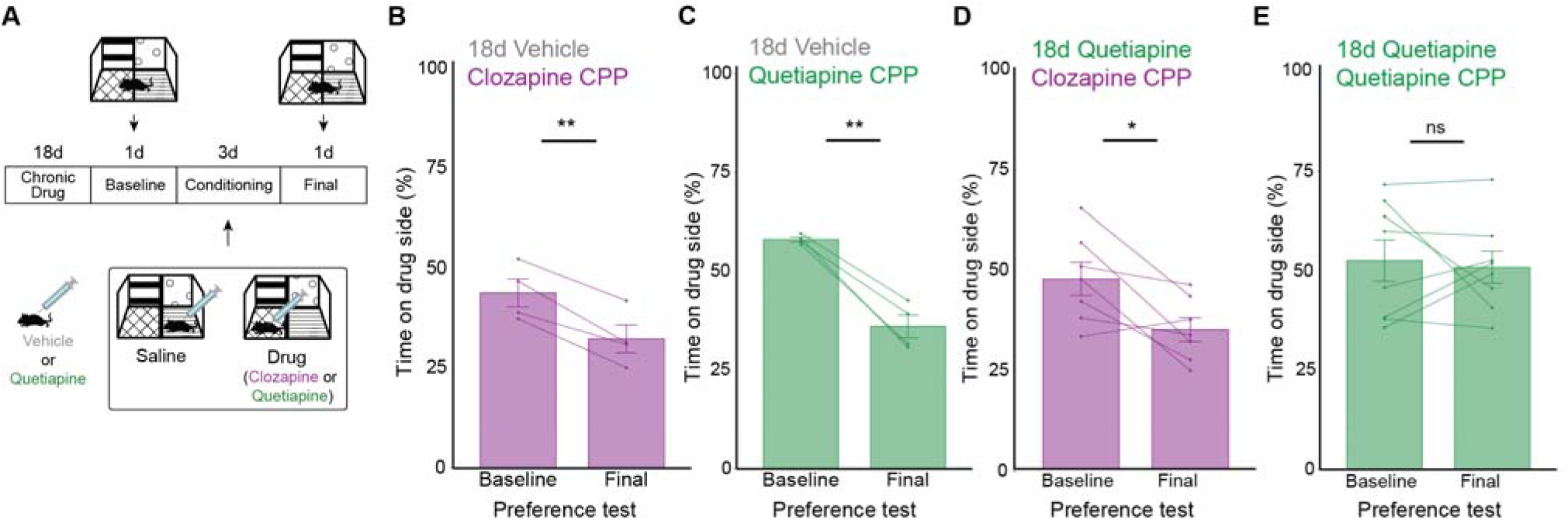
Chronic quetiapine injections do not cause cross-tolerance to clozapine conditioned place aversion. A) Timeline of conditioned place experiment. Baseline Preference Test and Final Preference Test take place when mice are off drug. Conditioning period includes am saline treatment in one context and pm, drug treatment in the other context. B) Mice chronically treated with vehicle (4% DMSO) avoid the clozapine (4 mg/kg) side after conditioning (N=4 mice, Student’s Paired t-test, **p=0.005, t=7.423). C) Mice chronically treated with vehicle (4% DMSO) avoid quetiapine (15 mg/kg) side (N=4 mice, Student’s Paired t-test, **p=0.003, t=8.456). D) Mice chronically treated with quetiapine (15 mg/kg) avoid the clozapine (4 mg/kg) side after conditioning (N=7 mice, Student’s Paired t-test, *p=0.025, t=2.980). E) Mice chronically treated with quetiapine (15 mg/kg) neither prefer nor avoid the quetiapine (15 mg/kg) side (N=8 mice, Student’s Paired t-test, ns, p=0.757, t=0.322). All error bars represent SEM.

**Figure S3:**
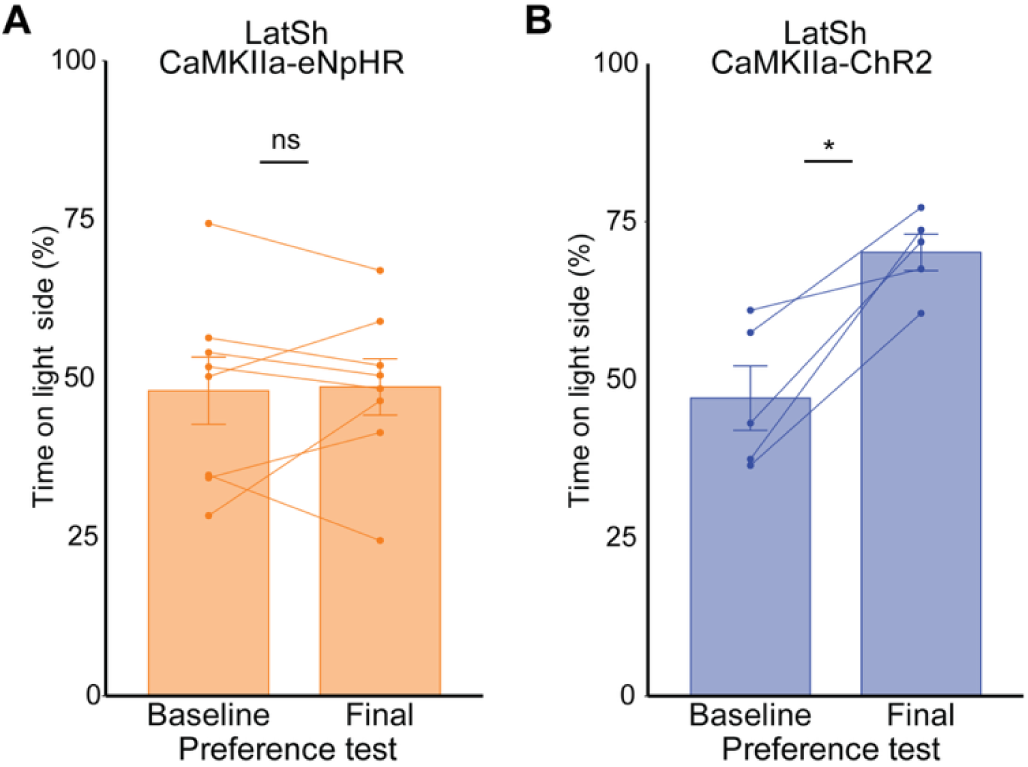
Nonspecific inhibition or excitation of LatSh NAc neurons does not drive Real Time Place Aversion in mice. A) WT mice expressing CaMKIIa-eNpHR in the LatSh NAc neither prefer nor avoid the orange light side (N=8 mice, Students Paired t-test, ns p=0.866, t=-0.175). B) WT mice expressing CaMKIIa-ChR2 in the LatSh NAc prefer the blue light side (N=5 mice, Students Paired t-test, *p=0.010, t=-4.660).

**Figure S4:**
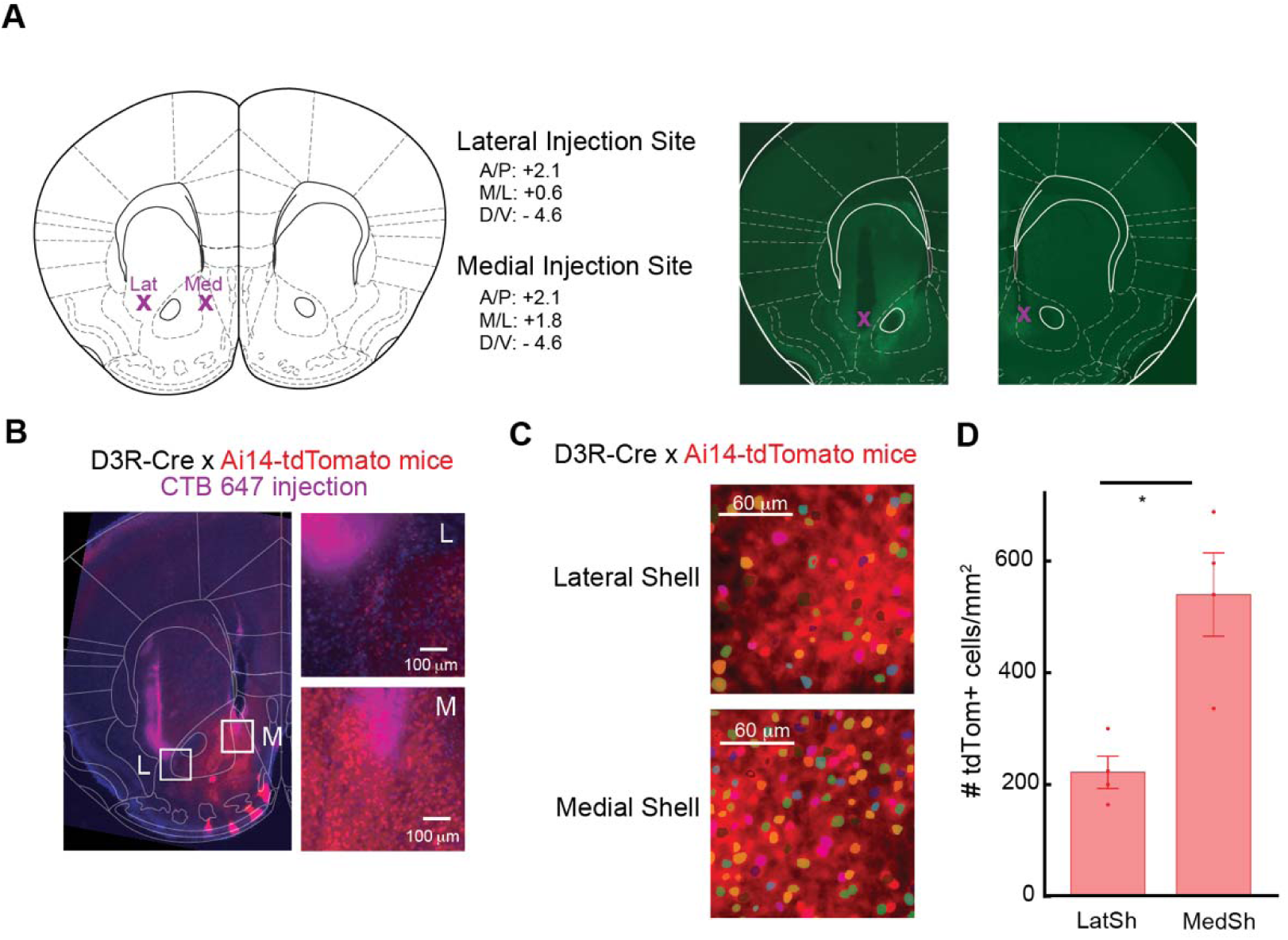
D3R neurons are more concentrated in MedSh NAc compared to LatSh NAc. A) Approximate injection sites and fiber locations are shown in coronal sections. Representative images of Cre-dependent GCaMP injections into LatSh NAc and MedSh NAc in D3-Cre mice show typical viral spread and fiber placement. B) CTB dye injections show ability to target MedSh and LatSh NAc separately. C) Representative images and cell masks used to quantify D3R-expressing in MedSh and LatSh NAc. Masks were drawn using Cellpose software with manual verification. See methods section for more detail. D) Quantification of D3R (tdTomato) cells in MedSh NAc compared to LatSh NAc (N=4 FOVs, Welch unpaired T-test, *p=0.017, t=-3.98). Error bars represent SEM

**Figure S5:**
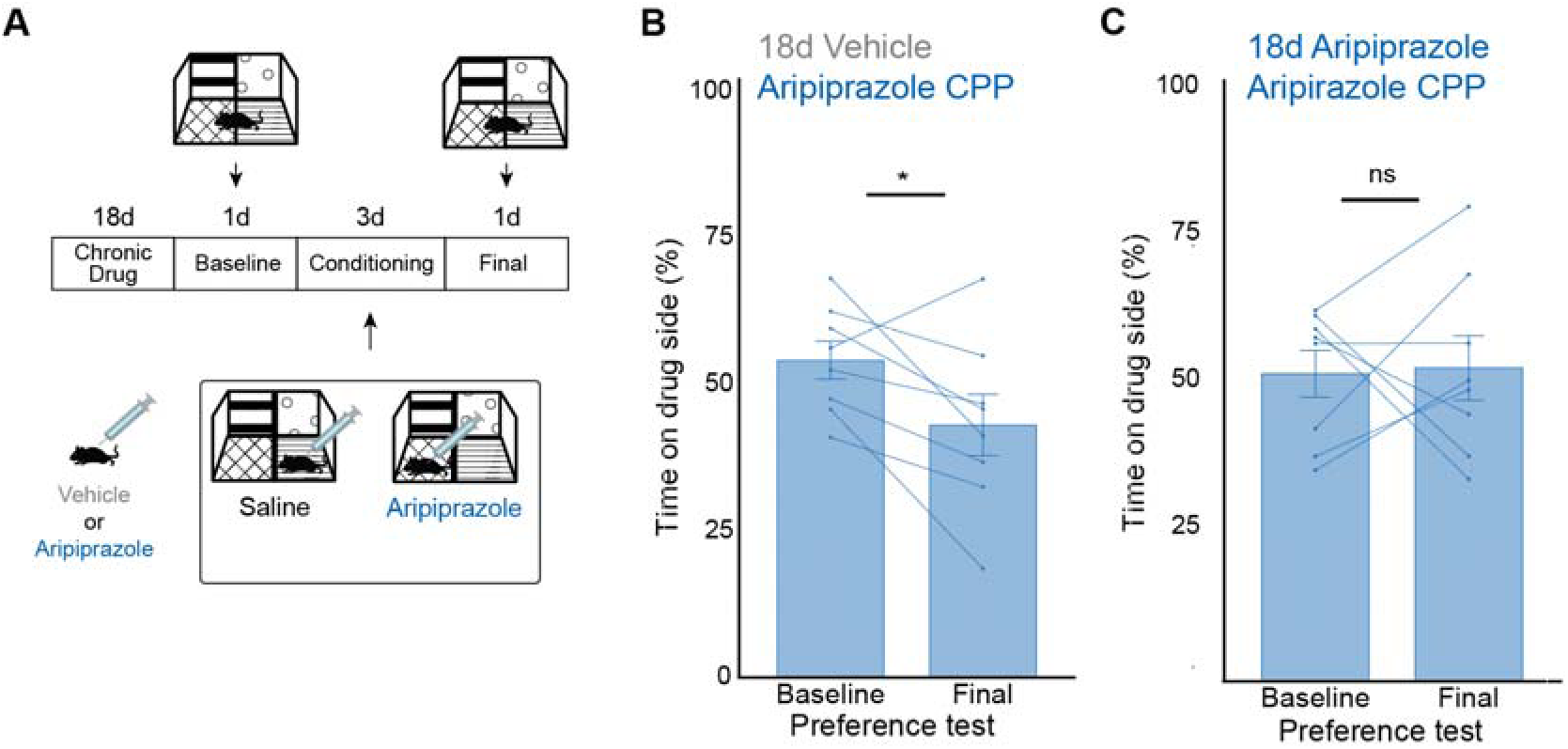
Mice become tolerant to the aversive effects of aripiprazole. A) Timeline of conditioned place experiment. Baseline Preference Test and Final Preference Test take place when mice are off drug. Conditioning period includes am saline treatment in one context and pm, drug treatment in the other context. B) Mice chronically treated with vehicle (4% DMSO) avoid the aripiprazole (15 mg/kg) side after conditioning (N=8 mice, Student’s Paired t-test, *p=0.040, t=2.519). C) Mice chronically treated with aripiprazole (15 mg/kg) neither prefer nor avoid the side paired with aripiprazole (15 mg/kg; N=8 mice, Student’s Paired t-test, ns, p=0.100, t=-0.145). All error bars represent SEM.

